# Accurate processing of ultra-short immune signals by single macrophages

**DOI:** 10.1101/2025.10.26.684377

**Authors:** Gabriel Mercado-Vásquez, James V. Vizzard, Betul Celiker, Junjie Xia, Andrew G. Wang, Abinash Padhi, Savaş Tay

## Abstract

Accurately interpreting short-lived signals is challenging within noisy cellular microenvironments. It is not known if cells can distinguish different signaling molecules under transient exposure. Here, we explored the temporal limits of signal detection by single macrophages. Microfluidic experiments monitoring NF-κB dynamics showed that macrophages strongly respond even to 1-second pulses of cytokine and pathogen ligands. Information theory showed that short-pulse response is highly specific to the stimulating ligand, comparable to that under long-term stimulation. Macrophages were mainly sensitive to the duration of cytokines, and the dose of pathogen ligands. Stimulus duration altered the ranking of response strengths among different pathogenic ligands. Mathematical modeling showed that receptor cooperativity is crucial for robust responses to transient signals, while receptor pathway variability leads to accurate signaling in fluctuating environments. These findings demonstrate that dynamic transcription factor specificity is preserved across varying signal durations and uncover network-level mechanisms that accurately distinguish transiently encountered threats.

## Introduction

Cellular processing of dynamic immune signals requires balancing sensitivity, response speed, and tolerance to benign stimuli. Cells detect and process a wide range of external signals during immune response. The cellular microenvironment is complex and presents dynamic signals that vary across multiple time scales. Innate immune cells like macrophages recognize pathogen associated molecular patterns (PAMPs) to detect the presence of pathogens (*1*–*3*) and secrete cytokines such as tumor necrosis factor alpha (TNF), interferons (IFNs) or interleukins (ILs) (*4*–*6*) to relay this information to other cells. Changes in the type, dose, and time-course of these signaling molecules indicate potential threats to surrounding cells and tissues (*7*). However, due to the noisy nature of the cellular environment as well as dynamic protein secretion by individual cells, the concentration of signaling molecules fluctuate across a wide range, complicating their accurate detection (*8*–*10*). Rapid and precise sensing of signaling molecules is crucial to contain infections and avoid tissue damage. On the other hand, delayed or improper responses can prevent pathogen clearance or resolution of the initial response, increasing the risk to the organism (*6*). Understanding how cells accurately process complex and variable signals—and identifying the conditions under which this processing fails—is critical for advancing our knowledge of immune regulation, and for accurately modeling and controlling infection and inflammation.

Intrinsic noise due to fluctuations in transcription or protein interactions as well as pre-existing cellular variability can inhibit the sensitivity of cells and change their overall response to external signals (*11*–*14*). Biological noise is even more relevant in scenarios where the stimulating ligand is present at low doses or for extremely short periods of time (*12*, *14*). The signaling inputs received by macrophages can be very short due to changing ligand concentration profiles across space and time (*10*, *15*–*18*). Processing of short-duration signals poses a fundamental challenge for all sensing systems, whether physical or biological in nature. It is unknown if individual immune cells can respond to and accurately process chemical signals with extremely short duration. For most of the cells in the immune system, it is still unclear how small of a signal they can detect, and in which context ligand detection will trigger a signal-specific transcriptional response, and when specificity fails.

Cells encode extracellular signal information into the dynamics of transcriptional pathways (*6*, *19*–*22*) like NF-κB, which is then decoded into specific gene expression programs on a population and single-cell level (*23*–*28*). In addition to the signaling molecule type, the response to external stimuli also depends on the characteristics of the input signal, such as its duration, magnitude and frequency, which convey critical information about the extracellular environment. The temporal profile of the input signal modulates transcription factor dynamics, gene expression outcomes and response variability in single cells (*20*, *21*, *25*, *29*, *30*). Responding specifically to dynamic signals is crucial for properly fighting infections and preventing autoimmunity. Single-cell imaging studies showed that primary macrophages specifically respond to persistent (long-duration) stimulation of pathogen signals, creating distinct NF-κB transcription factor dynamics that tightly depend on the ligand type (*19*, *26*, *27*). These results showed that macrophages accurately encode and decode chemical signals that are persistent and long term. However, little is known about how quickly and accurately macrophages respond to transient signals via NF-κB or other transcriptional pathways, and whether the specificity of their signaling response is conserved under transient (short-pulsed) stimulation. Overall, the temporal limits of pathogen and cytokine sensing by individual immune cells is unclear, and is not known if cells retain their ability to distinguish between different ligand types under a short, transient exposure.

To address these questions, we investigated the limits of sensing in individual primary macrophages exposed to short chemical pulses. Our experimental and mathematical studies asked how quickly and efficiently immune cells detect a signal perturbation in their environment and whether they adapt their response in the presence of fluctuating stimuli. Through microfluidic live cell stimulation experiments and single-cell tracking of NF-κB transcription factor dynamics, we investigated quantitative responses under small and transient perturbations, and compared them to behavior under persistent (hours-long) stimulation. We experimentally tested how cells respond to temporally variable signals that target both extracellular and intracellular receptors, stimulating primary macrophages with short pulses lasting 1, 10, 100 and 1000 seconds with cytokine and pathogen-derived signals. These experiments tested how fast the macrophages can respond, and whether initial exposure is sufficient to create a full NF-κB response in each cell. We also investigated the differences in response characteristics for longer durations vs. very short exposure.

We find that primary macrophages effectively respond to extremely short perturbations of cytokines and pathogen derived signals that last for a few seconds, and process the information encoded in the characteristics of these signals with high specificity, similar to long-term persistent stimulation. Depending on the stimulated pathway and the characteristics of the signal (amplitude and duration), macrophages exhibit high plasticity, quickly tuning their response according to changes in ligand dose and duration. Our experiments revealed two modes of sensing in stimulated macrophages: for pathogenic molecules their response is stronger with shorter exposure but higher doses of ligand, whereas for cytokine detection the response increases for lower doses but longer exposures. We then built a mathematical model that integrates the interactions between receptors and signaling molecules and the transduction of extracellular signals to activate NF-κB. Our model recapitulates the NF-κB activation under signals with variable amplitude-durations for different receptor pathways. We generalized our results to any receptor pathway activating NF-κB and propose a parameter space that defines the receptor activation modes. Finally, we mathematically study the effects of extrinsic and intrinsic noise in individual macrophage response using our model. Overall, our study explores the physical limits of signal sensing in macrophages using the NF-ΚB system as a model and elucidates the mechanisms that enable cells to respond sensitively and rapidly to environmental signals.

## Results

To achieve short-pulsed chemical stimulation and measure the capacity of primary murine macrophages to process transient signals, we used a microfluidic cell culture and stimulation device and tracked NF-κB dynamics in individual cells. This automated system allowed us to rapidly modulate the concentration and temporal profile of signaling molecules and control the duration of the cellular stimulation (*31*). The microfluidic system also controls environmental conditions to viably culture primary bone-marrow-derived macrophages (BMDM) obtained from transgenic mice expressing the fluorescently labeled p65/RelA transcription factor (mVenus-RelA) (*19*), and enables time-lapse fluorescent microscopy to monitor NF-κB pathway activation in individual cells.

To validate fast chemical delivery to cultured cells, we first calibrated our system by flowing short bursts of fluorescent dye to microfluidic culture chambers and monitoring the fluorescent signal with microscopy. Fast chemical signals with a duration as short as one second can be achieved within the microfluidic chambers (Supplemental Figure 1A, and Supplemental Video 1). We also performed control experiments by flowing short bursts of fresh culture media to detect any background response induced by shear-stress or by the rapid perturbation of the cell environment. We did not find any detectable flow-induced response in NF-κB nuclear localization (Supplemental Figure 1B). This system delivers chemical stimulation pulses as short as 1 second to live cells, with 6% variability in timing across all analyzed cells (coefficient of variation ∼ 0.06).

### Individual macrophages detect extremely short pulses of pathogen signals

Using our system, we stimulated individual macrophages with various NF-κB ligands, tracked nuclear translocation of the NF-κB transcription factor p65 in cells, and analyzed its dynamics using custom image processing software (Figure 1A-B). For each stimulation condition that we tested, we analyzed more than 200 individual cells over the course of 4 hours.

**Figure 1.**
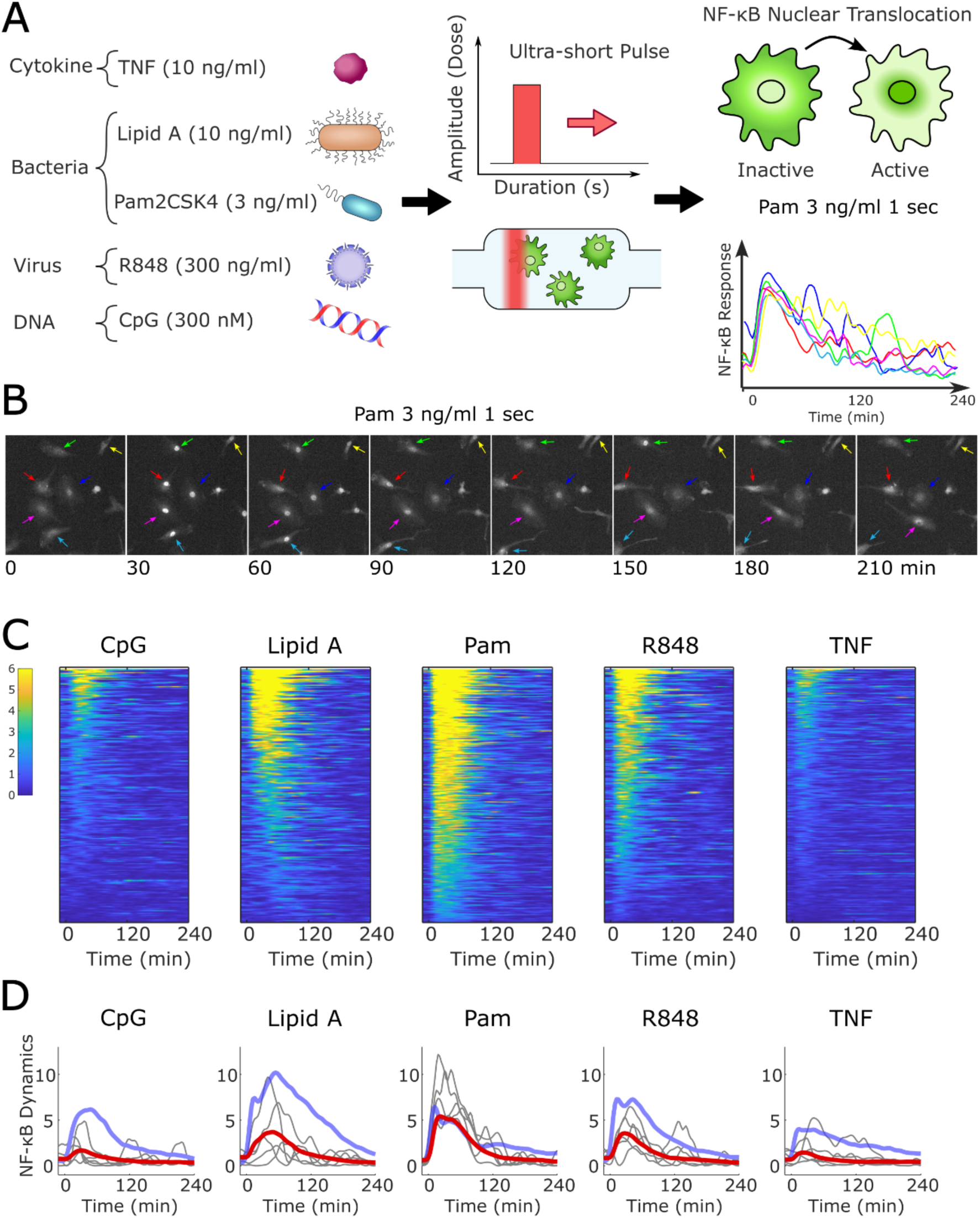
Single macrophages respond to ultra-short pulses of cytokines and pathogen derived ligands, and population variability depend on signal type: A) Using microfluidic culture combined with fluorescence microscopy, the dynamics of the NF-κB transcription factor p65 was recorded in primary bone marrow derived macrophages stimulated with short pulses of various ligands; 10 ng/ml TNF, 10 ng/ml Lipid A, 3 ng/ml Pam2CSK4, 300 ng/ml R848 and 300 nM CpG. The cells were exposed to a traveling chemical pulse of 1-second duration, by flowing the ligand loaded medium over cultured cells. B) Example of primary macrophages stimulated with a 1-second pulse of Pam2CSK4. Colored arrows indicate the cells whose traces are depicted in A-right. C) Single-cell heatmaps of the nuclear translocation of the transcription factor NF-κB in response to one second stimulation of ligand. Each row represents a single-cell trace, x-axis is time. D) Red line represents the mean NF-κB activation within the population under one second stimulation for different signaling molecules. Blue line represents the mean NF-κB activation within the population for the constant stimulation in Supplemental Figure 1C. Thin lines are NF-κB traces from 5 single cells randomly chosen in each case.

We tested the response of macrophages to short pulses of approximately one second duration with five different types of stimuli, including one cytokine and four pathogen associated molecules at relevant concentrations. To simulate intracellular communication, we stimulated the cells with the proinflammatory cytokine TNF at a concentration of 10 ng/ml. For pathogenic stimulation we targeted 4 different receptors of the TLR family: Lipid A at 10 ng/ml and Pam2CSK4 at 3 ng/ml are analogs of bacterial molecules which bind to Toll-like receptors (TLR) 4 and 2, respectively. For viral and foreign DNA exposure, we used the ligand R848 at 300 ng/ml and the single stranded CpG DNA dinucleotide at 300 nM, whose main receptors are TLR7 and the TLR9, respectively. The doses we used induced persistent NF-κB activation in primary macrophages when supplied in a continuous manner in previous studies (*19*, *32*). With these molecules, single macrophages exhibit high response specificity under persistent stimulation, creating NF-κB dynamics that are tightly determined by the stimulating ligand type (*19*, *26*, *32*, *33*) (Figure 1A).

Our experiments revealed that single macrophages respond in a sensitive manner to short pulses. Under each ligand stimulation, many cells produced strong NF-κB localization peaks (Figure 1B-D, and Supplemental Video 2). Several individual macrophages effectively detected the transient signals of cytokine and pathogen-associated molecules that last as short as one second. This signal duration is in striking contrast to many transcriptional and translational processes that govern the NF-κB pathway and many other biological processes, which happen on time-scales ranging from tens of minutes to several hours. The response time of NF-κB activation is in the tens-of-minutes time scale, and an input time scale of 1 second is therefore hundreds to thousands of times faster compared to the typical activation timescales of NF-κB observed in single cells.

We also found that the detection range as well as the strength (amplitude) of NF-κB nuclear translocation was ligand dependent, as shown by the different p65 activation dynamics exhibited for each ligand type (Figure 1C-D). Single-cell analysis revealed significant variability within the macrophage population in response to short pulses, and the extent of single-cell variability depended on the type of ligand used. Overall, these experiments showed that single macrophages can detect extremely short exposures to many self-secreted and foreign signaling molecules, and strongly activate the NF-κB pathway in response.

### NF-κB activation is transient in individual macrophages exposed to short signaling pulses

Modulation of the NF-κB dynamics leads to different transcriptional outcomes (*19*–*21*). The population mean of NF-κB dynamics in our short-pulsed stimulation experiments showed that macrophage response occurs during the first 1 or 2 hours after ligand exposure and quickly dampens. This shows a relatively shorter activity compared to the dynamics of macrophages in the presence of persistent stimulation (Figure 1D, Supplemental Figure 1C); NF-κB activity under persistent (constant) stimulation shows a much longer activity duration, typically lasting 10 hours or more (*16*, *19*, *20*, *34*).

The type of genes regulated by NF-κB has a strong correlation with the duration of nuclear occupancy and chromatin accessibility (*27*, *28*). Fast nuclear translocation allows transcription of genes that are accessible at early times, whereas a longer nuclear occupancy induces expression of late genes (*28*). At the population level, our experiments indicate that ultra-short stimulation elicits relatively short NF-κB dynamics that last for up to two hours, suggesting that most of the cells direct their response towards the rapid activation of early genes specific to the stimulated pathway, which are typically involved in the production of negative feedback proteins that help inactivate the NF-κB pathway.

### Single-cell analysis reveals subpopulations of early, late and non-responder macrophage profiles under short-pulsed stimulation

The response of the cells is quickly attenuated under short pulses at the population level. Analysis of single-cell traces, however, reveals a small fraction of individual cells that exhibit long-term NF-κB dynamics lasting for the entire course of the experiment. Even in the case of TNF stimulation, which elicits the weakest response, some cells displayed persistent NF-κB dynamics. This variability cannot be explained with the 7% input signal dose variation (AUC, CV=0.07) observed in the microfluidic chamber (Supplemental Figure 1A). To better understand the response of individual cells stimulated with short pulses of ligands, we analyze their dynamics by computing 8 different signal features extracted from the NF-κB traces (*19*) (see Figure 2A). As single-cell NF-κB dynamics transfer external signal information into gene expression outcomes, detailed analysis of dynamic NF-κB profiles enables better understanding of how this information transfer occurs (*16*, *19*, *25*, *27*, *30*). The dynamic NF-κB features we analyzed include area under the curve (AUC) at early (<45 min) and late (>45 min) time points, indicating the total nuclear activity of RelA, time to activation, amplitude of peak activation, time to reach peak activation, the oscillatory dynamics, the speed of translocation, and duration of the nuclear NF-κB activity.

**Figure 2.**
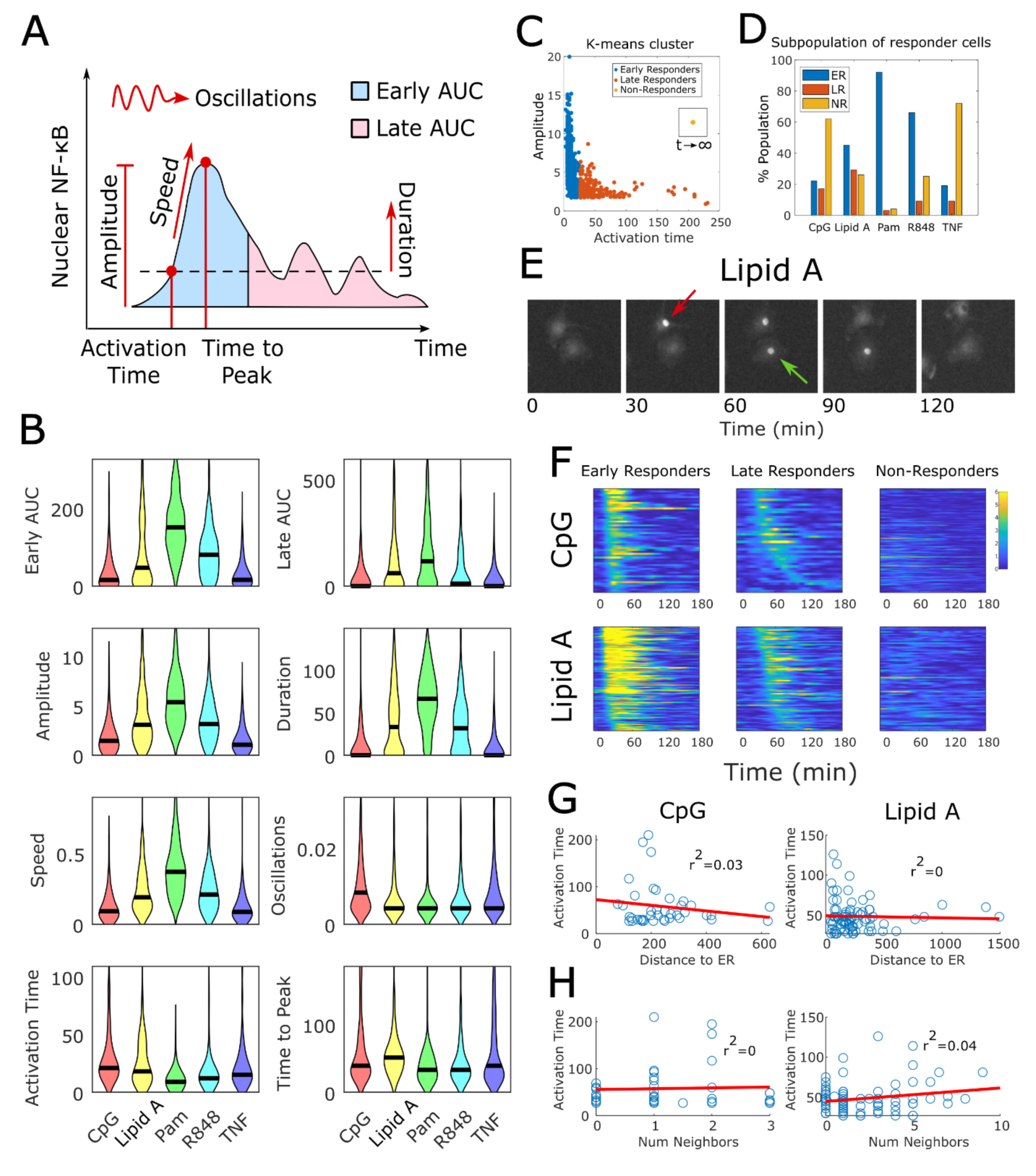
Single-cell analysis reveals sub-populations of early, late and non-responding macrophages to short pulsed stimulation. A) Schematic of the 8 features extracted from the NF-κB traces: the area under the curve (AUC) at two time points, early (<45 min) and late (>45 min), activation speed, maximum activation (peak), duration of the activation (time above activation threshold), oscillatory dynamics (first harmonic), time to activation and time to peak. B) Violin plots for the comparison of the dynamic features for each ligand. Black line represents median. C) k-means clustering of the macrophage’s response based on the activation time and amplitude of each trace. For illustration purposes, we included the non-responder cells that never reach the minimal activation level in a separate frame. E) Example of late responders for Lipid A stimulation. The red arrow points toward the early responders whereas the green arrow corresponds to the late responder. F) Heatmap of the Early, Late and Non-Responders. G) Activation time vs Distance to the nearest Early Responder (ER). H) Activation time vs Number of Early responders (neighbors) around a late responder.

Figure 2B shows the violin plots of the 8 dynamical features. Pam2CSK4 stimulation displays the strongest response, with the late and early AUC, amplitude, speed and duration showing the highest values. For the oscillatory behavior however, CpG and TNF were the ligands that induced oscillations with higher frequencies. This contrasts with previous studies with non-specialized immune cells, where the oscillatory dynamic is attenuated or completely terminated once the ligand has been removed (*18*, *24*).

In most of the dynamic features, Pam exhibits the strongest response, contrary to previous persistent simulation studies that found Lipid A as the most potent immunostimulant (*22*). Interestingly, when we measure NF-κB traces under persistent stimulation (Figure 1D, Supplemental Figure 1C), Lipid A indeed emerges as the most potent ligand. This comparison shows that the strength of cellular response to pathogenic inputs strongly depends on the duration of stimulation, and not only on the identity of the ligand itself.

Quantification of the dynamic features shows that the variability of the NF-κB dynamics is ligand dependent. The early AUC and amplitude (Figure 2B) show that a significant fraction of cells stimulated with Lipid A exhibits a very high sensitivity, reaching amplitudes higher than the average response in Pam stimulation. However, the duration of the transcription factor in the nucleus shows a heterogenous response, with some cells exhibiting no response at all. Furthermore, cells treated with CpG or Lipid A exhibited a broader range of activation time, with some cells initiating NF-κB activation much later than those stimulated with TNF or R848. Interestingly, Lipid A induces the largest time to peak in average (> 50 min), indicating a late response of cells stimulated with this ligand (Figure 2F).

We then asked whether this shift in the time response corresponds to a population behavior or if it is due to heterogeneity at the single-cell level. To answer this question, we perform a k-means clustering of all the NF-κB traces based on their activation time and amplitude to reveal distinct subpopulations of cells with different dynamic response patterns (Figure 2C, see Methods). We found three subpopulations: 1) a subpopulation of early responders that exhibits a fast response (<25 min) with high amplitude; 2) a subpopulation of late responders (>20 min) with low amplitude; and 3) a subpopulation of non-responder cells that never reach the minimum activation threshold. As indicated in Figure 2D, these three subpopulations are present across all five ligands that we tested, nevertheless, their proportions vary depending on the ligand. Pam induces the lowest fraction of late responders (∼3%), whereas Lipid A displays the largest population of late responders (∼30%). As shown in Figure 2E-F, cells become activated even one hour after stimulation with Lipid A.

As stimulation with pathogenic signals can induce cytokine secretion and paracrine signaling, we asked whether this late response is due to detection of secondary paracrine signals, rather than the primary stimulus. To answer this question, we analyze the correlation between the activation time in late responders and their physical proximity to early responders (see Methods). If paracrine secretion is mediating the late response, the cells near early responders are more likely to quickly activate. As shown in Figure 2G-H (Supplemental Figure 2A), we did not find any correlation between the activation time and the distance to the nearest early responder nor to the number of early responders in a certain vicinity around the late responders. Additionally, we measured the correlation of the NF-κB dynamic features between the late and the nearest early responders without finding any strong correlation (Supplemental Figure 2B). These results indicate that the late response is most likely to be driven by factors intrinsic to the NF-κB pathway rather than external cell-to-cell communication.

### Macrophages distinguish ultra-short pathogen signals and respond specifically

Cells accurately discriminate between different ligands under persistent stimulation and activate NF-κB in ligand-specific dynamical patterns (*19*, *27*, *32*, *34*, *35*). However, processing of short signals is challenging for sensing systems compared to long term signals, as the sensor may not have sufficient time to integrate signals and distinguish them from others and from background noise. Short signal exposure could pose significant challenges to specificity in NF-κB, leading to indistinguishability between pathogenic, proinflammatory and anti-inflammatory signals in physiological settings. Such break-down of inflammatory signal specificity may lead to insufficient pathogen clearance or tissue damage (*6*, *36*). Overall, it is not known if cells retain their ability to distinguish between different ligand types under a short, transient exposure.

Since macrophages can detect extremely short perturbations in the environment, we proceeded to determine how accurately they can process the information carried by such short signals. As macrophages continuously encode information from the extracellular environment and decode it into a dynamic NF-κB trace (*32*, *33*)(Figure 3A), we can use an information theoretical approach to analyze how well a macrophage is able to discriminate between different ligands. This framework has been previously used to quantify the specificity of immune response in the presence of persistent pathogen associated signals (*14*, *19*, *26*, *31*). We start by calculating the mutual information (MI) of the NF-κB traces between two different input ligands. In this context, the pairwise MI is a measure of the amount of information that the NF-κB traces contain about the stimulating source. Comparing the response between two different inputs allows determining how specific the response is relative to each other. If the NF-κB traces for two different ligands are completely distinguishable, the mutual information would contain 1 bit of information, whereas complete indistinguishability will be equal to zero. Furthermore, as the NF-κB traces are dynamic, this analysis would determine at which time point the responses are more distinguishable between each other.

**Figure 3.**
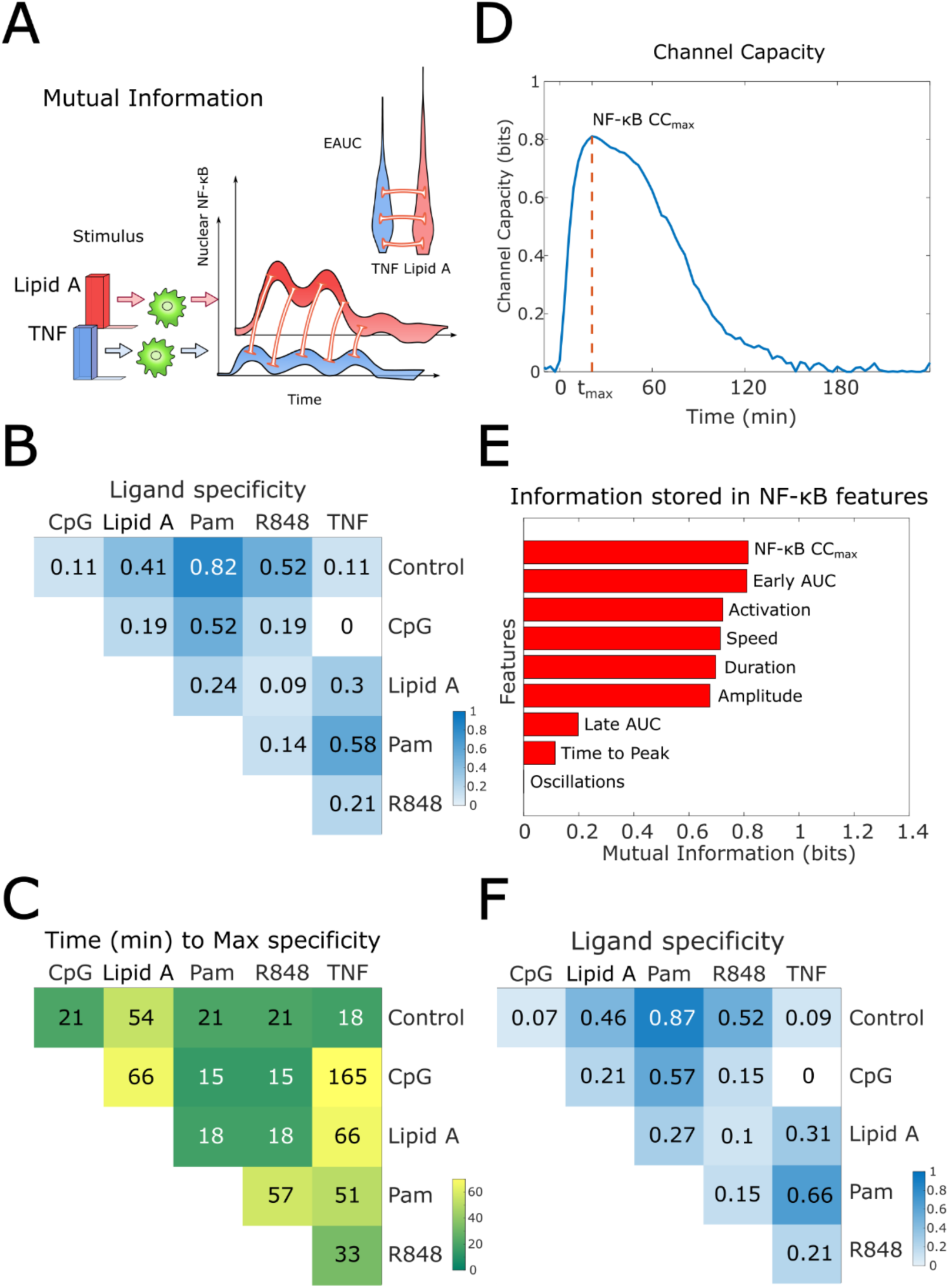
Macrophages exhibit a high response specificity and accurately distinguish signals with ultra-short stimulation: A) Schematic of mutual information. Macrophages responses for two different inputs are compared at each time point. The same comparison is made with the distribution of NF-κB features. B) Pair-wise measurement of the maximal mutual information of the NF-κB traces for each pair of ligands. Mutual information of 1 indicates high specificity or distinguishability in the response. C) Time at which the maximum specificity is reached in B). D) Channel capacity (CC) of the NF-κB traces for different ligands as a function of time. At time *t*_*max*_the CC reaches its maximum. E) Channel capacity based on the NF-κB features. F) Combining dynamic features with the NF-κB signal at *t*_*max*_ we calculate the mutual information for a pair of ligands. This shows the specificity in the macrophage response to each ligand traces. Stimulation to Pam shows almost perfect distinguishability.

Figure 3B shows the maximum pairwise MI between two inputs, whereas Figure 3C indicates the corresponding time where this maximum is reached. MI is highest when comparing any stimulated group with the control group. Stimulation with Pam gives the maximal MI close to complete distinguishability, followed by R848. On the other hand, stimulation with Lipid A shows a lower distinguishability that occurs at a later timepoint due to activation of late responders. These differences indicate how in response to a quick pulse of cytokines or pathogenic ligands, macrophages process the information at different time scales depending on the stimulating source. When comparing the specificity between ligands, the highest distinguishability that we found was between Pam and TNF, followed by Pam and CpG. Interestingly, CpG and TNF show the lowest MI, indicating that macrophages do not distinguish between these two ligands for short stimulation.

Channel capacity (CC) represents the maximal flux of information that a system can transmit. It is defined as the maximal mutual information between two variables, here the NF-κB dynamics and the stimulation source. In this context, the channel capacity measures how well cells distinguish between multiple input sources. We computed the channel capacity with the raw NF-κB traces at different time points (Figure 3D) and found a maximal channel capacity of *CC*_*max*_ = 0.82. This is close to the time in which NF-κB reaches maximal nuclear translocation upon Pam stimulation, which gives the strongest response. Surprisingly, the channel capacity for fast pulsed stimulation is comparable with that observed for primary macrophages under persistent stimulation (Supplemental Figure 3A), where the maximal channel capacity is *CC*_*max*_ = 0.92. Similar values have been reported for immune cells in the context of hours-long persistent stimulation (*14*, *19*, *37*). This result surprisingly indicates that primary macrophages respond to short signals nearly as effectively as they would under constant stimulation conditions. Furthermore, even when the duration of the signal is of the order of seconds, the cell response and its capacity to distinguish between different inputs can last for more than one hour. However, for the persistent stimulation case the distinguishability is maintained for much longer times (Supplemental Figure 3A).

Including additional dynamic NF-κB features improves response specificity depending on the ligand origin (*33*). We calculated the channel capacity for eight additional dynamic features. As shown in Figure 3E, among the most informative features are the early AUC, the amplitude at the time point *t*_*max*_ (*NF* − *κB CC*_*max*_), and the speed of activation. Therefore, the biggest differences in the response to short-time perturbations with cytokines and pathogenic ligands are related to the early activity of the transcription factor p65 and its nuclear translocation speed. Combining these most informative features, we calculated the mutual information between different ligands. As shown in Figure 3F, we observe an increase of mutual information, with the cells being able to discriminate between control and Pam2CSK4 with MI=0.87. This value is much closer to channel capacity under persistent stimulation, which represents an almost perfect distinguishability of different ligands. These results indicate that macrophages maximize their signal distinguishing ability by storing information in multiple dynamic features of the NF-κB nuclear translocation.

### NF-κB dynamics encodes both the duration and amplitude of pathogen and cytokine signals

To elucidate how macrophages process information about the amplitude and duration of signaling inputs of various durations, we expose primary macrophages to different ligand profiles using our microfluidic platform with pulses of duration 1, 10, 100 and 1000 seconds (Figure 4A). To better understand the effect of signal duration vs amplitude, we keep the number of molecules delivered to cells constant across all stimulation profiles by decreasing the ligand dose by a factor inversely proportional to the duration. We chose three different stimulants to model different processes during proinflammatory signaling: TNF-TNFR models cell-cell communication, Pam2CSK4 and the TLR2 models extracellular sensing of bacteria, and R848 binds to TLR7 located in the endosome to model endosomal receptor mediated detection of pathogenic signals.

**Figure 4.**
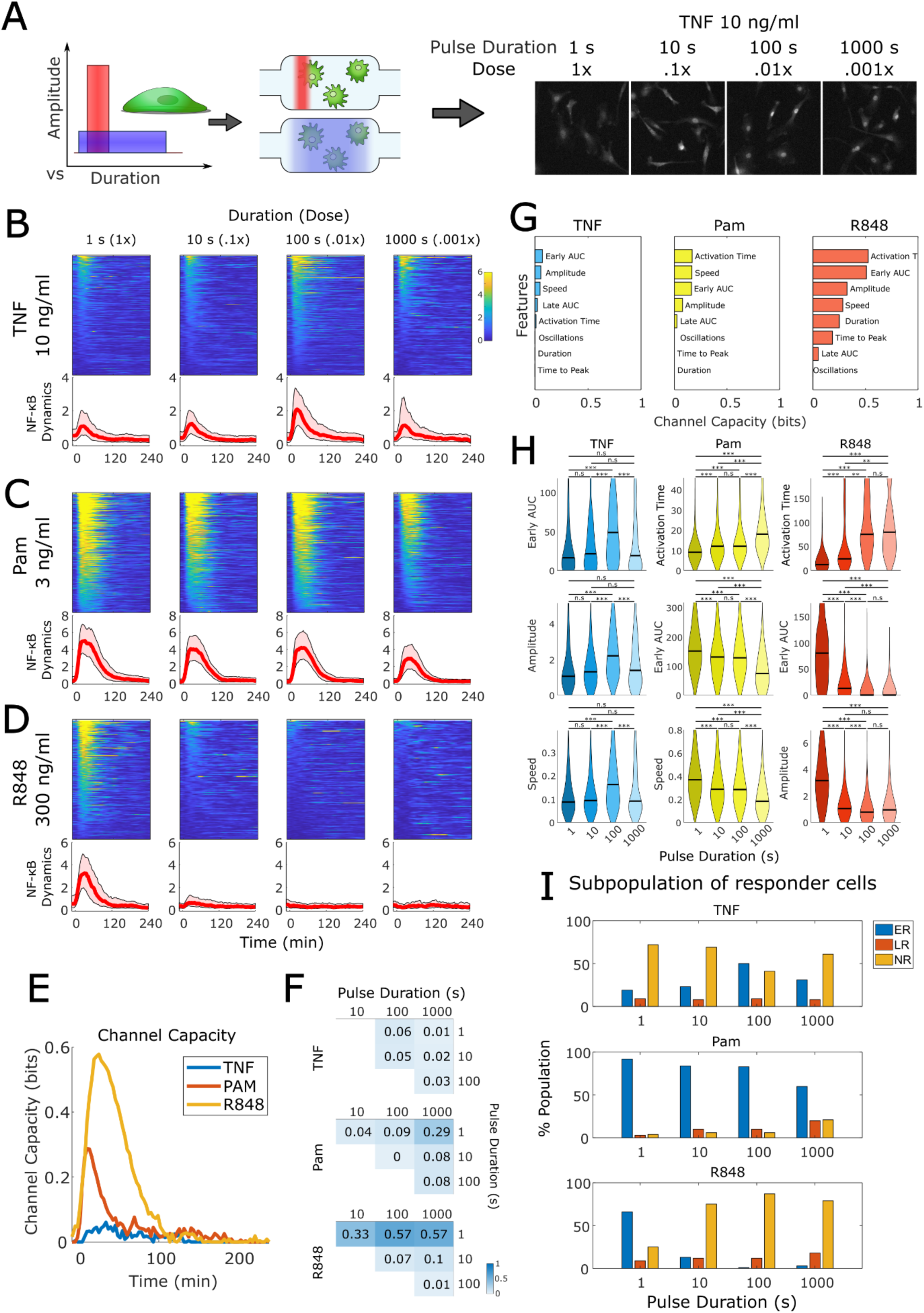
Macrophage activation decodes duration and amplitude of stimulation: A) Primary macrophages are stimulated with the cytokine TNF and the pathogenic ligands Pam2CSK4 and R848 with pulses lasting for 1, 10, 100 and 1000 seconds. In each case, the area of stimulation (dose x duration) is kept constant. B-D) heatmap and traces of the nuclear NF-κB response for different durations and doses for TNF, Pam and R848. Light color areas represent the 25^th^ and the 75^th^ percentile, whereas the thick red line represents the median. E) Channel capacity in response to different durations and amplitudes of the same ligand. F) MI comparing the distinguishability to input duration and amplitude for TNF, Pam and R848. G) Which features of the NF-κB dynamics better decodes information of the input shape. H) Distribution of the most informative features in G. Black line represents median values. I) subpopulation of early, late and non-responders for each ligand and pulse duration.

Our experiments showed that macrophages integrate information of the shape of the signal input and modify their response accordingly. For macrophages exposed to the same stimulation area (concentration multiplied by duration), NF-κB exhibits distinct dynamics depending on the duration of the input, its concentration and its identity, whether it is a cytokine or a pathogenic signal. For TNF stimulation (Figure 4B, and Supplemental Video 3), we observed non-monotonic behavior. The number of activated cells and strength of activation increased with longer stimulation, but only up until 100s. At the population level, a 100-second pulse of TNF doubles the NF-κB amplitude compared to a 1-second pulse. However, low concentrations can diminish the macrophage’s response, even with longer stimulation (i.e. 1000-second pulse).

On the other hand, stimulation with the lipopeptide Pam2CSK4 induced a subtle decrease in the NF-κB activity when increasing the duration of the signal (Figure 4C, and Supplemental Video 4), in this case, higher doses induce higher amplitude response. Surprisingly, for the small molecule R848 which specifically targets the intracellular receptor TLR7, longer exposures but lower doses elicited a minimal to null response (Figure 4D, and Supplemental Video 5). This all-or-nothing dynamic is characteristic of non-immune cells such as fibroblast or human cancer cells in the context of cytokine stimulation (*16*, *28*, *38*). However, as has been reported, the response of macrophages to pathogen ligands may appear analog, with a gradual but continuous increase of the number of activated cells and their activation strength (*19*, *39*, *40*). This could suggest that macrophages adjust their response depending on the type of signaling molecule from bacterial components, RNA, virus or cytokine.

### Macrophages are sensitive to the dose of pathogenic signals and the duration of cytokines

To determine how effectively macrophages decode the identity and shape of the input signal, we calculated the channel capacity of NF-κB across different pulse durations and amplitudes in response to a single ligand (Figure 4E). In this case, the channel capacity measures how well the cells can distinguish between different input shapes for a specific ligand when the stimulation area is fixed. As expected, the response to R848 exhibits the largest channel capacity, which reflects the fact that lower doses fail to trigger NF-κB response. On the other hand, although at the population level the differences in TNF stimulation are clear, CC shows a very low value, not capturing the differences in single-cell response. This is due to the large subpopulation of non-responders that dominates the distribution in the calculation of the CC. Conversely, stimulation with the lipopeptide Pam2CSK4 shows a relatively high channel capacity within a very short time window after simulation (<20 min), indicated that the biggest differences in response with different input durations and amplitudes take place in the early response.

Additionally, we computed the pairwise MI to compare the capacity of the cells to distinguish between two pairs of inputs with different duration and amplitudes (Figure 4F). Overall, these results show that, at short time scales, there are significant differences in the individual macrophage’s response as a function of the amplitude and duration of the stimulus, suggesting a potential differential transcriptional response depending on the characteristics of the stimulus shape.

Since macrophages adapt their response to input duration and amplitude, we asked which features of the NF-κB traces better capture the adaptation process at the single-cell level. We computed channel capacity of the NF-κB dynamic features for each ligand, which elucidates how macrophages store information of the input shape in the NF-κB features. As shown in Figure 4G, the early AUC, speed, amplitude and activation time are the features that show the highest channel capacity overall, which is also prominent from the distribution of these features (Figure 4H). Interestingly, the NF-κB activation time becomes more relevant in the distinguishability of pathogenic signals than for the cytokine TNF. Overall, the most informative features are related to the early dynamics of NF-κB. The early nuclear activity better decodes information about the input shape for transient perturbation. Differences in the nuclear occupancy of the transcription factor NF-κB in response to stimulating signal have been related to changes at the transcriptomic level and epigenetic reprograming (*20*, *21*, *25*, *27*, *41*), suggesting that changes in pulse duration and dose could lead to changes in cell state.

Finally, we asked whether the percentage of late responders remains the same across different input durations for a given ligand. We performed a k-means clustering of the amplitude and activation time centered on the same points as for the one second stimulation case (Figure 4I). For TNF, the fraction of early responders reaches maximum at the 100-second pulse, whereas the fraction of late responders remains the same for all durations. Conversely, for Pam stimulation the subpopulation of late responders increases with pulse duration, whereas the percentage of early responders significantly decreases. For R848 we can see the switch between early responders to non-responders due to the decrease in ligand dose.

Overall, our experiments demonstrated that macrophages respond to signaling molecules in two different ways: for endogenous signals like cytokines, macrophages primarily respond to signal duration, with longer stimulation inducing stronger response when the signal is above a minimal threshold. In contrast, macrophage predominantly responds to signal amplitudes for pathogenic signals, activating stronger and faster when the dose of the stimulating ligand increases, even with short-lived stimuli.

### Mathematical modeling reveals signal-dependent transitions between high- and low-dose filtering in receptor pathways

Although stimulation with the cytokine TNF and the pathogenic signals Pam2CSK4 and R848 converge into the activation of IKK module and activate NF-κB, the mechanisms regulating these pathways upstream the IKK are different. Using mathematical modelling, we investigated which components of the pathway can explain the characteristic dynamics observed in the response of macrophages to ultra-fast signals, and whether intrinsic and extrinsic noise plays a relevant role in processing these signals.

Several mathematical models have been developed to simulate the dynamics of NF-κB activation (*15*, *17*, *21*, *25*, *31*, *42*, *43*), some of which incorporate ligand-receptor engagement (*12*, *18*, *19*, *28*, *30*). In this work, we developed a simple mathematical model (Figure 5A) that captures NF-κB response to brief stimulations. The model incorporates ligand–receptor interactions and the subsequent signal transduction to the IKK complex in a minimal yet representative manner, using minimal parameterization while preserving the key properties intrinsic to the stimulating pathway. Importantly, our model uses a Hill Function formalism to represent receptor interactions, where *K* and the coefficient *n* of the activation function (Hill function) that transduce the signal from the active receptors to the IKK complex. This activation function summarizes the multiple protein-protein interactions that occur upstream IKK (See Methods).

**Figure 5.**
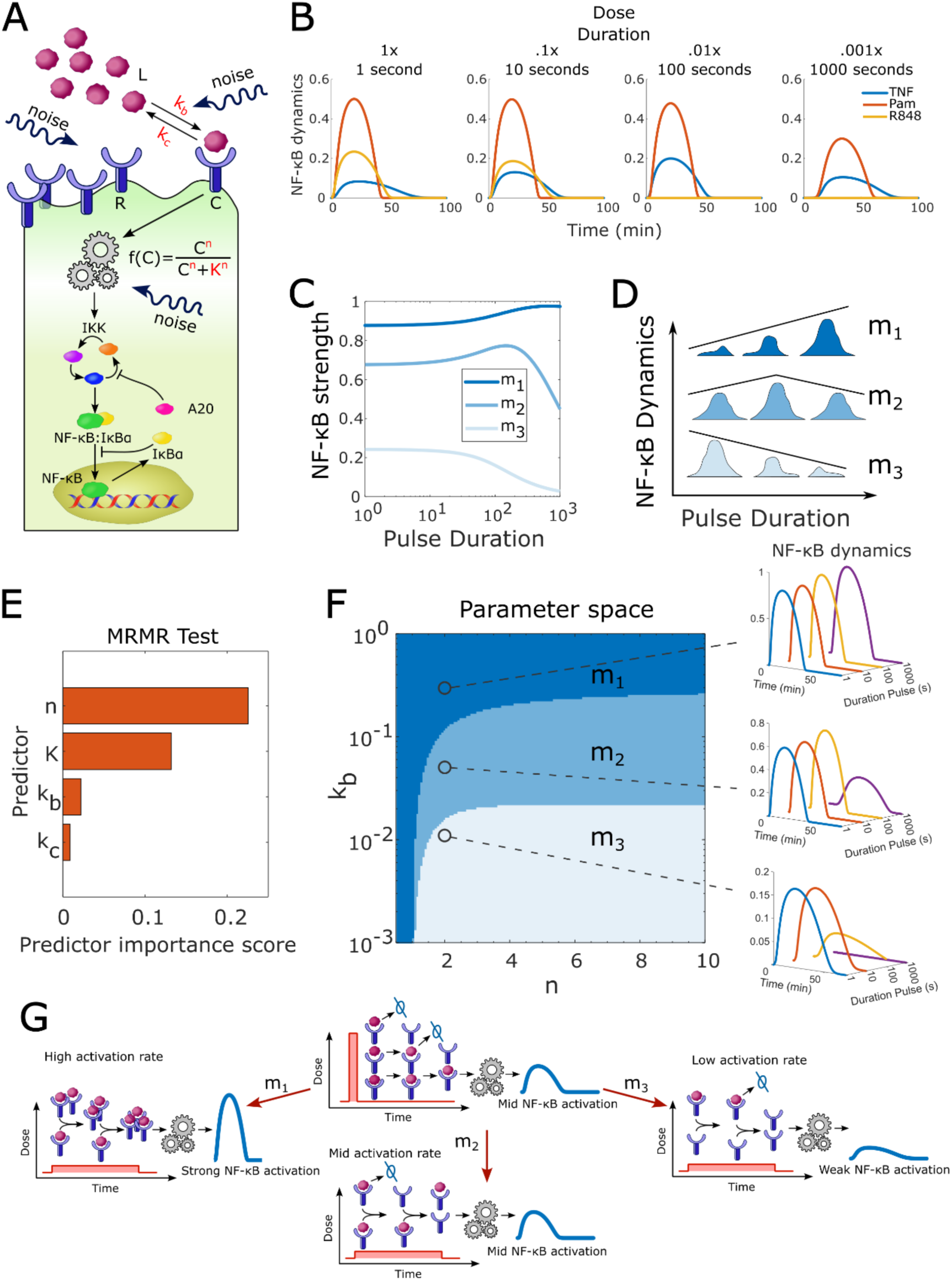
Mathematical model predicts three distinct activation modes in cells: A) Schematic of our simplified mathematical model. Extracellular ligands interact with the receptors of the cell. The external signal is transduced by a mechanism intrinsic to each receptor pathway and converges into the activation of the IKK complex, which activates NF-κB. B) This model recapitulates the three different activation patterns observed in response to TNF, Pam and the small molecule R848. C) Our model predicts three modes of activation with variable input duration. D) Schematic of the three main modes in which primary cells activate in response to extracellular signals with longer (lower) pulse duration (amplitude): in mode *m*_1_ the activation peak monotonically increases; in *m*_2_ the activation peak is non-monotonous; and in *m*_3_ the activation peak monotonically decreases. E) Minimum redundancy maximum relevance (MRMR) test that identifies the most relevant parameters that predicts the NF-κB activation mode. F) Phase diagram in the parameter space where the activation peak exhibits one of the three activation modes. Our model predicts phase transitions between the three activation modes as a function of the free parameters. We show representative traces for the response to four pulses with different durations with the following fixed values: from bottom to top *k*_*b*_ = .01, .05, .5 1/*s*, the rest of parameters are *k*_*c*_ = .03 1/*s*, *K* = .01 and *n* = 2. G) Conceptual figure of the mechanisms that lead to the three activation modes as described in the main text.

First, we tested whether the simple model reproduces cell activation in response to short signals, and whether it can capture distinct dynamic modes depending on the pulse duration. Parametrization of our model via global optimization allowed us to find the best parameters that fit the population dynamics for different pulse durations (see Table 2). Our model recapitulated the distinct dynamics exhibited by three stimulated pathways (Figure 5B). Consistent with literature (*44*–*46*), our model predicted higher activation rates for TNFR and TLR2 than for TLR7 (see Table 2). The Hill coefficient *n* displays the lowest value (*n* ≈ 1.2) for TNFR, whereas for TLR2 and 7 we found *n* ≈ 1.8 and *n* = 10, respectively. Interestingly, similar values of the Hill coefficient have been found for TLR4 under constant stimulation (*12*).

To better understand the principles that govern different dynamic modes, and to know which model components better explain NF-κB activation with variable dose and pulse duration, we varied model parameters across a wide range of values. To make this process even more general, we rewrote our equations, expressing the receptor activation rate in units of the maximal ligand dose, and normalizing the number of active receptors to the total receptor expression (see Methods). This step allows us to generalize our results to any receptor pathway and stimulation dose. This general model reproduced not only the two dynamic modes for a broad range of parameter values but also predicted a third mode in which the activation strength monotonically increases with duration pulse (Figure 5C). We then classified the dynamics of the NF-κB strength for the different pulse durations into three categories (Figure 5D): *m*_1_ (monotonic increase), *m*_2_ (non-monotonic) and *m*_3_ (monotonic decrease). Using the minimum redundancy maximum relevance (MRMR) algorithm (Matlab, see Methods), we identified the Hill coefficient *n* as the parameter with the highest MRMR score (Figure 5E), meaning that cooperativity between the active receptors is a better predictor of the NF-κB modes in response to fluctuating stimulation. Surprisingly, our model predicted phase transitions between the three dynamic modes when increasing *k*_*b*_ for different values of *n*, while keeping *K* and *k*_*c*_ constant (Figure 5F and Supplemental Figure 4).

Our results showed that differences in the activation dynamics are due to an interplay between how fast receptors activate (*k*_*b*_ limited), how many of them can be simultaneously active (*k*_*c*_ limited), the minimal number of active receptors to trigger response (*K* limited), and whether they exhibit positive or negative cooperativity (*n* limited) (Figure 5G). For receptor pathways exhibiting positive cooperativity (*n* > 1), simultaneous activation of multiple receptors amplifies the transduced signal. When receptor activation rate is high, even minimal doses of ligand can trigger NF-κB response. In this case, longer stimulation pulses allow for the accumulation of active receptors, further enhancing NF-κB activation strength (*m*_1_). This effect becomes more pronounced when receptor deactivates rapidly, and only long-lasting signals will be transduced.

At intermediate receptor activation rates, only ligand doses above a threshold effectively transduce the signal, whereas very low doses elicit weak or negligible NF-κB activation (*m*_2_). For low receptor activation rates, only high ligand doses can activate NF-κB, whereas low doses have minimal effects on activation, even when the signal duration is extended (*m*_3_). Conversely, for receptor pathways exhibiting negative cooperativity (*n* < 1), simultaneous activation of multiple receptors can diminish overall activation strength. In this scenario, fewer simultaneously active receptors contribute more effectively to total activation when the signal persists (*m*_1_).

In summary, pathways with high cooperativity but low activation rate will most likely operate as high-dose filters (*m*_3_), triggering strong responses only in the case of high doses. On the other hand, pathways with poor or negative cooperativity but high activation rates will function as low-pass filters (*m*_1_), letting only low signals to fully activate the pathway. Interestingly, the TNF receptor lies right on the border between these two modes with *n* close to 1 and with a high activation rate, whereas TLR 2 and 7 show low activation rates with high cooperativity.

### Intrinsic and extrinsic factors modulate NF-κB dynamics to transient perturbations

For transient signals, variation of the number of stimulating molecules surrounding a single cell are likely to affect the overall response. Although our microfluidic cell culture device robustly performs fast chemical stimulation, there is some local variation in ligand dose. As shown in Supplemental Figure 1A, for fast stimulation, the variation of the total amount of signaling molecules reaching a single cell is 7% around the mean. However, the MRMR analysis (Figure 5E) showed that variation of the parameter *k*_*b*_ (which expressed the effective dose a single cell is exposed to) is not one of the most relevant parameters to predict the existence of the different dynamic modes in NF-κB. Instead, the cooperativity (*n*) and the minimal number of active receptors to reach half-maximal activation (*K*) are more impactful for this outcome.

Variability of the receptor levels is thought as one of the main sources of extrinsic noise responsible for the heterogeneity of the innate immune cell response (*12*, *28*, *29*, *33*, *47*–*49*). On the other hand, stochastic fluctuations in protein-protein interaction (intrinsic noise) also contribute to the variability in cell response (*11*, *12*, *14*, *21*, *29*). Using our mathematical model, we investigated how variation of the receptor levels modulates the NF-κB response and its dynamic features at the single-cell level for cells stimulated with 4 pulses with different durations and doses as for our experimental pipeline (Figure 6A).

**Figure 6.**
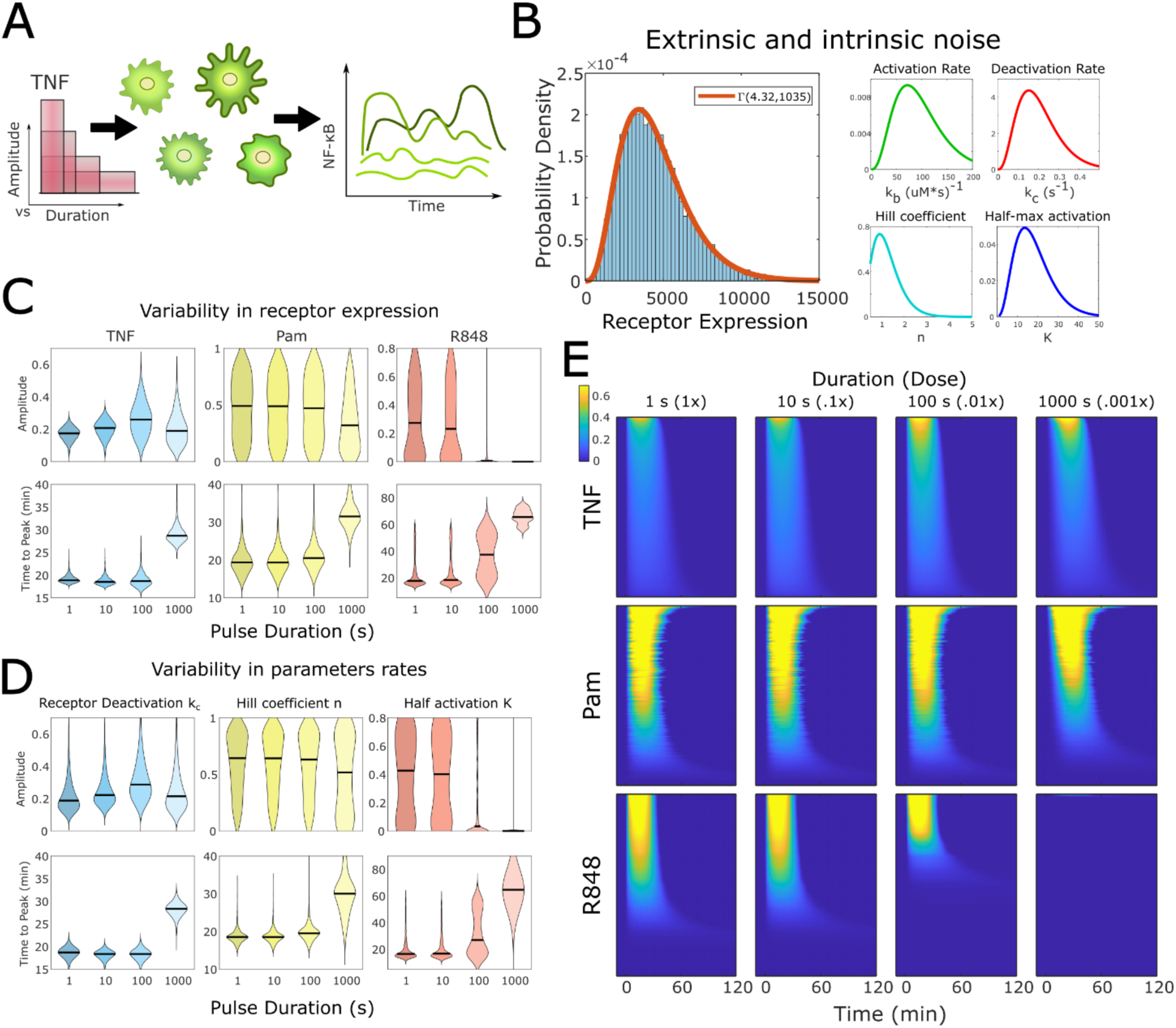
Extrinsic and intrinsic noise explain the variability in NF-κB dynamical features in single cells: A) For each condition we generate 1000 synthetic cells with variable parameters and receptor expression. Then we stimulated them following a similar pipeline as in Figure 4A. B) Measurement of the receptor levels fits a Gamma distribution, showing the variability of protein expression in single cells due to extrinsic noise. We proposed similar variability for the rest of the parameters in our model. C) Violin plots for the NF-κB activation amplitude and time to peak when the receptor levels are varied in our simulation. Each violin plot represents the distribution of 1000 cells. Black lines represent median values in the population. Variation of the receptor levels partially explains the observed variability in the experimental NF-κB dynamical features. D) This variability is better explained when more parameters are subject to stochasticity. For TNF stimulation we vary the receptor deactivation rate *k*_*c*_, whereas for Pam and R848 we vary the Hill coefficient *n* and the constant *K*, respectively. F) Heatmap of the NF-κB traces predicted by our model when both the receptor levels and the biochemical rates in D stochastically fluctuate.

We obtained the distribution of receptor expressions in macrophages for TNFR, TLR2 and TLR7 through flow cytometry measurements (Figure 6B). We fit the receptor expression to a Gamma distribution as it is usual for protein expression in cells (*13*). Variation of the receptor expression helps explain the variability of the activation amplitude for the three ligands (Figure 6C). However, single-cell receptor variability did not completely reproduce the behavior we observed experimentally. In our simulation of Pam stimulation, variation of the receptor levels could not explain variability in time to activation and the presence of early and late responders at longer pulse durations. For TNF, the NF-κB amplitude does not show the high variation as in the analysis of experimental data.

We hypothesized that these subtle differences may be related to the variation of other rates in our model. We performed a variation of the reaction rates using Gamma distributions centered at the optimal rates for each pathway (Figure 6B). This gave the distribution of NF-κB amplitudes and time to peak when independently varying each of the parameters in our model (Figure 6D, Supplemental Figure 5). Varying the Hill coefficient *n* in Pam stimulation generates a broader distribution of the time to peak, and it is the only parameter that produces a subpopulation of late and early responder cells (Figure 5F). For TNF, variation of *k*_*c*_ shows a broader range for the amplitude for the four doses. We then combine the variation of receptor expression with the variation of the Hill coefficient *n* for Pam stimulation to generate more realistic traces of the NF-κB activation (Figure 6E). Similarly, we reproduced NF-κB traces for TNF and R848 stimulation for variable receptor expression and fluctuating *k*_*c*_ and *K*, respectively.

Although variability of the receptor levels leads to heterogeneity in the NF-κB activation, it is not the only source of noise that affects the cell response in fluctuating environments. Other sources of noise such as variation of the kinetic rates at which proteins interact influence the activation dynamics in single cells. Our simulations showed that fluctuations of the receptor-receptor interaction (cooperativity) have a primary influence in the speed at which a signal is transduced. Likewise, stochasticity in the deactivation of receptors contributes to increasing or decreasing the activation strength in individual cells.

## Discussion

Using microfluidic live cell analysis and mathematical modeling, we investigated the limits of signal detection by single primary macrophages in response to cytokines and pathogen derived signaling molecules. We stimulated primary macrophages with physiologically relevant doses of several ligands at short durations in the 1 second scale while monitoring the dynamics of the NF-κB transcription factor p65, one of the main transcription factors involved in the inflammatory response in infection, autoimmunity and cancer.

Our studies revealed that macrophages can quickly recognize and respond accordingly to pulses of a stimulating ligand that last for as brief as one second, exhibiting a highly specific response. Surprisingly, the specificity under short pulses is comparable with that observed under sustained stimulation (*19*, *32*). When compared to fibroblasts stimulated with low doses for a prolonged duration, primary macrophages outperformed their specificity, exhibiting a much higher distinguishability between multiple ligands even when the signal is brief (*37*). This finding highlights the powerful sensing capabilities of specialized first-responder cells like macrophages. Even though macrophages are exposed to a lower number of signaling molecules under short stimulation, their capability to respond to external cues and their capacity to trigger differential responses remain high. The fact that a minimal perturbation in the environment triggers a highly specific response indicates that macrophages are equipped with highly sensitive and sophisticated machinery for the detection and sensing of extracellular signals, and are able to detect and aggressively respond to any minimal changes of signaling molecules in their surroundings. This remarkable ability to rapidly and accurately respond to environmental cues opens new questions in our understanding of how immune cells sense fluctuating signals and program their respond to short-lasting stimulus.

The response to transient signals can differ substantially from that observed in persistent stimulation. In epithelial cells, minute-long pulses of the cytokines IL-1b and TNF elicit stronger responses than continuous exposure (*50*), even if the cells are exposed to more signaling molecules in total. In neuroblastoma cells, 5 minutes exposure to TNF initiates transcription of only early genes (*20*), whereas late genes require longer TNF exposure. Likewise, our experiments suggest that the response of macrophages in the presence of small perturbations is mostly dedicated to turn off their own activity, as opposed to the case of persistent stimulation, where the cells display a longer response and lead to upregulation of many genes regulating cell fate decisions.

Our experiments show that primary macrophages employ different mechanisms to sort transient signals depending on the input origin. In intercellular communication through the cytokine TNF, macrophages employ a mechanism of detection that filters extremely short and transient cytokine inputs, even if the strength of the stimuli is high, in favor of long-lasting signals. This indicates a high degree of specialization in macrophages for robust communication in noisy environments. Persistent exposure to cytokines not only leads to longer and stronger activation (*24*, *50*), but also influences transcription, as many genes are only accessible for longer duration of the nuclear localization of the transcription factor NF-κB (*18*, *24*, *25*, *28*, *41*). Although less sensitive than primary macrophages, HeLa cells exhibit strong activation to 10 second pulse of 100 ng/ml TNF (*51*). Interestingly, the percentage of apoptotic cells after TNF treatment follows a non-monotonic behavior with the stimulus duration, with 1 minute stimulation driving same number of apoptotic cells than continuous exposure (*51*).

For the pathogenic signals Pam2CSK4 and R848, macrophages filter low amplitudes or doses, initiating activation only when the pathogen load is high enough, representing a real threat for the immune system. In RAW 264.7 macrophages, transcription of the NF-κB subunit RelA, along with a specific subset of target genes, occurs only under high-dose of LPS stimulation (*40*). In fibroblast, the fraction of activated cells increases with the dose of LPS ligand (*30*). Nevertheless, a non-monotonic increase of the NF-κB activation strength with the ligand dose has been also observed in TLR4 stimulation (*52*). This suggests that the mode of cell activation strongly depends on the specific properties of the stimulatory pathway, rather than representing a universal mechanism for pathogen detection.

We developed a simple mathematical model that integrated receptor-ligand interaction with signal transduction to activate NF-κB. Our model recapitulated the activation patterns observed in primary macrophages and predicted the existence of at least three dynamic modes in the NF-κB activation that depend on properties inherent to the stimulatory pathway. In these modes the NF-κB amplitude depended on the duration and amplitude of the pulse. Receptors with low activation rates will preferentially show a monotonic decrease in the activation peak for longer signal durations (lower signal doses). On the contrary, receptors with high activation rates can exhibit a non-monotonic peak increase. However, modulation of the NF-κB response strongly relied on the cooperativity between the active receptors. Whereas in our model TNFR exhibits low cooperativity, the Toll-like receptors are more likely to cooperate between them. In most of the TLR pathways, formation of complex Myddossome structures is required to initiate NF-κB activation (*3*), which help explain the higher rates we predict in our model.

Intrinsic and extrinsic noise contribute to the variability in cellular responses to stimulation(*11*, *14*). Whereas intrinsic noise depends on thermodynamic fluctuations that affect biochemical reactions inside the cell, extrinsic noise depends on variations in the amounts of cell components, such as protein or RNA expression. Using our mathematical model, we identified potential sources of extrinsic and intrinsic noise that can explain the variability in the response of macrophages to transient signals. The expression of receptors involved in the recognition of extracellular signals partially contributed to explaining such variability, however, a better explanation can be given when we include cell-to-cell variability of the kinetic rates at which the molecules in the regulatory pathway transduce the signals.

In summary, we showed that macrophages distinguish short signals specifically, and have identified potential mechanisms in which cells can sort transient environmental signals and respond according to the characteristics of the signal, such as its type, dose and duration. These findings help elucidate the mechanism that innate immune cells employ in sensing extracellular and intracellular signals. Our findings can have potential consequences in our understanding of the immune response and in the use and application of small molecules in therapy.

## Limitations of the study

This study reports the signal specific response of single macrophages in an *in vitro* setting. Future experiments using *in vivo* models may further improve significance of our results. While we tested a number of signaling molecules that represent key sources, the possible number of all signals received by macrophages *in vivo* is a lot larger. Future experiments with a larger library of signaling molecules would be necessary to completely map macrophage response specificity. Further, since we use laminar flow to stimulate the cells in our culture chambers, we expect an effect of the convective forces combining with diffusion in the transport of ligands into the receptors. This plays an important role in determining the kinetic rates of the ligand-binding processes for most cases. Therefore, the kinetic rates found here may vary from most of the experiments where diffusion is the main driving force involved in ligand-protein interaction. However, at physiological levels, we expect our analysis to be closer to what can be happening in a real environment where ligands spread out by non-diffusive transport through the bloodstream.

## Acknowledgements and funding information

This work is supported by NIH Grants R35GM148231. The authors would like to thank the University of Chicago Animal Resources, Center, and UCFlow for providing facilities and equipment. The mVenus-RelA (RelA^V/V^) C57BL/6J mice *(19)* was a gift from Alex Hoffmann laboratory.

## Author contributions

Conceptualization, S.T. and G.M.V.; Methodology, G.M.V.; Investigation, G.M.V., J.V., B.C., J.X. and A.P.; Writing – Original Draft, G.M.V.; Writing – Review & Editing, G.M.V., J.V., B.C., J.X., A.G.W. and S.T.; Funding Acquisition, S.T.; Resources, S.T.; Supervision, S.T.

## Methods

### Lead contact

Please direct further questions and requests for reagents and resources to the lead contact, Savas Tay

### Data and code availability

Code is uploaded and available at https://github.com/tay-lab/Accurate-processing-of-ultra-short-immune-signals-by-single-macrophages.

### Reagents

For cell differentiation we used murine m-CSF (Peprotech) reconstituted in ultrapure water and aliquoted and stored at -80C. For ligand stimulation we used the murine tumor necrosis factor alpha (TNFα) (R&D Systems), PAM2CSK4 (PAM) (Invivogen), ultrapure Lipid A from E. coli O111:B4 (Invivogen), ODN 1668 (CpG) (Invivogen), and R848 Resiquimod (Invivogen).

### Mouse models

We bred endogenously tagged mVenus-RelA (RelA^V/V^) C57BL/6J mice (*19*) under specific pathogen-free conditions at the University of Chicago. All experiments were conducted in accordance with the NIH Guide for the Care and Use of Laboratory Animals approved by the University of Chicago Institutional Animal Care and Use Committee.

### Cells

The BMDMs used in this study were extracted, differentiated and cultured as previously described (*32*). In short, we isolated bone marrow cells from the tibias and femurs of 6–12-week-old male and female RelA^V/V^ mice and resuspended in complete media supplemented with 20 ng/mL m-CSF. We harvested the BMDMs on days 7-9 and loaded and seeded in microfluidic culture chips in complete media.

### Microfluidic device fabrication

The fabrication of the microfluidic devices was according to our previously published protocol (*31*). Briefly, we used silicon wafer master molds to fabricate multilayer polydimethylsiloxane (PDMS) in a 10 to 1 ratio of PDMS (Momentive) monomer and catalyst. This mix was poured or spin-coated to form the control and fluid layer, respectively. We bonded to the fluid layer to the control layer using oxygen plasma followed by overnight baking. We punched control and flow inlets, followed by glass bonding.

### Microfluidics enabled live-cell imaging

Live-cell imaging was performed with our setup previously described elsewhere. Similarly, the details of our microfluidic live-cell stimulation and imaging setup have been previously described elsewhere (*53*). In short, using a GUI customized in MATLAB we electronically controlled pneumatic valves that were connected to the microfluidic device inlets to control on-chip valves. An in-situ microbiome box was used to maintain the right temperature, humidity and *CO*_2_ conditions for cell culture. Cells were harvested and seeded at a density of 10^7^ cells/ml into the device chambers previously incubated with 0.2 mg/mL fibronectin solution (Sigma-Aldrich) in PBS. After seeding, the cells were left unperturbed for 4-6 hours until they were attached and equilibrate in the microfluidic chambers. Then, the cells were stained for nuclear tracking with 1 μM Hoechst 33342 (Invitrogen) in phenol red-free RPMI (Gibco) media according to the producer instructions. Using a Nikon Ti2 epifluorescence microscope we acquired images of the mVenus-RelA (508-nm) and Hoechst (395-nm) at 6 min rate for 6 hours at 20x magnification. The images were recorded using a Hamamatsu, ORCA-Flash4.0 V2 camera. We did not observe any photobleaching or phototoxicity.

### Microfluidic stimulation

The reagents previously described were taken from -80 C stock and diluted in complete media. To preserve their integrity, the samples were stored on ice over the course of the experiment and delivered into the microfluidic chambers using polyetheretherketone tubing (VICI) at an input pressure of 4 psi.

### Quantification and Statistical Analysis

#### Image analysis and trace processing

For image analysis and cell tracking we used customized software in MATLAB as described in previous publication (*32*). Briefly, raw images acquired with the Nikon software were converted into MATLAB format and pre-processed for wide-field and background correction. Hoechst images were used to segment nuclei and track the cells over the entire course of the experiment. From the mVenus fluorescence images we extracted information of the NF-κB signal inside and outside the nucleus. We computed the ratio of nuclear/cytosol NF-κB. Death cells, dividing cells as well as cells out-of-focus or with any other imaging artifacts were removed to achieve a >99% accuracy in tracking (*32*).

#### Quantification of the NF-κB dynamic features

All the NF-κB features were calculated using MATLAB 2024a: for the early and late AUC we computed the trapezoidal approximation (trapz) at the intervals 0 to 45 min and 45 to 240 min, respectively. The amplitude and time to peak were calculated using the in-built function findpeaks. The speed was computed as the maximal rate change of the NF-κB levels in time using the in-built function diff in matlab. To compute the activation time and the duration of the activation we first computed the activation threshold, which is defined as 2 s.d. of the NF-κB level above the mean of the nuclear to cytoplasm ratio for unstimulated cells. The activation time is then defined as the first passage time of the NF-κB level to the activation threshold. The duration of activation is defined as the time the NF-κB level is above the activation threshold. Computing the first harmonic in the fast Fourier transform of the NF-κB traces we obtained oscillatory behavior in single cells.

#### Spatio-temporal analysis

With the activation time and amplitude values described in the above section, we performed a k-means clustering in MATLAB, performing 10 random replicates of the centroids and using Euclidean distance. This splits the data into two populations, the early and late responders. The third population of non-responders correspond to the cells that never reached the activation threshold and have no value for the activation time. Next, with the x and y-positions that we got from our cell tracking algorithm, we computed the nearest early responder neighbors around each of the late responder cells using Euclidean distance. We then compute the linear correlation between the NF-κB features of the nearest neighbors and the late responder cells.

#### Mutual Information

The mutual information of the NF-κB traces and their dynamic features was computed as described in previous work and in references elsewhere (*14*, *19*, *32*). Briefly, the mutual information *I* defined as

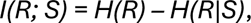

calculates the difference between the non-conditional Shannon entropy *H(R)* of observing a response *R* and the conditional entropy *H(R|S)* of observing the response given the stimulus *S*.

One can interpret mutual information as a metric that quantifies the amount of certainty in observing a response given that the cells were stimulated with a certain stimulus. If by looking at the response one can perfectly tell which stimulus was used, *I* will max out 1, otherwise it is lower than 1.

#### Minimum Redundancy Maximum Relevance (MRMR)

To understand which of the four parameters in our model could better explain the existence of the NF-κB response mode, we simulated more than 100,000 synthetic cells with their parameters randomly generated from a uniform distribution within reasonable boundaries, making sure that the values of TNFR, TLR2 and TLR7 fit from the experimental data fell into the parameter space. Then, we stimulated each individual cell with 4 pulses with constant stimulation area (similar to our experimental pipeline) and classified the NF-κB response according to the three groups *m*_1_, *m*_2_ and *m*_3_. Then, we apply the fscmrmr function in Matlab to rank the four predictors in our model for the classification of the NF-κB mode.

#### Mathematical Modelling

Our model describes the out of equilibrium interaction between free receptors (R) and ligand molecules (L), which leads to the formation of active receptors (C). An activation function (Hill function) summarizes the multiple interactions downstream the active receptors and upstream the IKK complex. We then used a minimal model inspired from (*42*, *43*) that captures the main behavior of the NF-κB activation, namely, the translocation of the transcription factor upon stimulation and the negative feedback given by IkBa and A20 proteins (see Jensen et al, 2012 for more details).

Stimulation of macrophage receptors activates a variety of ubiquination and kinase cascades which culminate in activation of the IKK complex that phosphorylates the inhibitor IkBa for its immediate degradation, releasing the cytoplasmic p65. Once free, the p65 rapidly imports into the nucleus and transcribes target genes and its own inhibitor IkBa. Newly synthesized IkBa sequesters NF-κB, translocating it back to the cytoplasm. If the stimulating signal persists, the whole process will be repeated leading to oscillation. This process results in a dynamic signal that characterizes the response of macrophages to stimulation.

Although stimulation of the TLR and TNFR converges into the activation of the IKK complex, and the subsequent activation of the transcription factor NF-κB, each receptor family has a unique regulatory pathway upstream IKK. The signaling of TNFR is mediated by the recruitment of the adaptor molecule preligand assembly domain (PLAD) (*54*). This molecule facilitates the formation of the TNFR trimer (*49*, *54*, *55*), which recruits the adaptor molecule TNFR-associated death domain (TRADD) within the cytoplasm. The adaptor molecule TNFR-associated factor (TRAF2) will then mediate the activation of IKK, finally leading to the activation of the NF-κB protein (*49*, *55*). On the other hand, stimulation with the diacylated lipopeptide Pam2CSK4 induces the formation of the stable heterodimer TLR2/6 with an m-shape that allows activation of the intracellular TIR domains (*2*, *56*). Although the adaptor protein myeloid differentiation primary response gene 88 (MyD88) is the main responsible for the activation of the IKK module that triggers the NF-κB response (*56*, *57*), TLR2 requires TIRAP/Mal for bridging to MyD88. Stable heterodimerization of TLR2 and TLR6 is achieved by the strong hydrogen bonds that interact with the peptide chains of the ligand and the exposure of surface residues of both receptors, therefore, the diacylated lipopeptide serves as a bridge for the receptor dimerization (*56*). Similarly, R848 binding induces dimerization of TLR7. This complex is required for the interaction with the intracellular TIR domain that communicates with MyD88. As mentioned before, MyD88 induces activation of IKK-NF-κB circuit (*1*, *2*, *44*, *56*, *57*).

Eq. (1) describes the interaction between the free receptors (M-C) and the ligand L, where M is the total amount of receptors in the cell. With rate *k*_*b*_ the receptor is activated due to the presence of the ligand, whereas with rate *k*_*d*_ the complex receptor-ligand is deactivated. In Eq. (2), the IKK complex in the neutral state (*IKK*_*T*_ − *IKK*_*a*_ − *IKK*_*i*_) is activated by a Hill function depending on the active complexes C, at rate at rate *k*_*a*_. In Eq. (3), active *IKK*_*a*_ is deactivated with rate *k*_*i*_ and transitions back to neutral form with rate *k*_*p*_.

In Eq. (4) active IKK induces activation of cytosolic NF-κB (*N*_*T*_ − *NFκB*_*n*_), which is then translocated into the nucleus at rate *k*_*in*_. This process is inhibited by cytosolic *IkBα*. Nuclear NF-κB is transported back to the cytosol due to the newly synthetized *IkBα* at rate *k*_*out*_. Dimeric NF-κB in the nucleus transcribes new mRNA molecules (Eq. 5) which are then translated into new *IkBα* (Eq. 6).

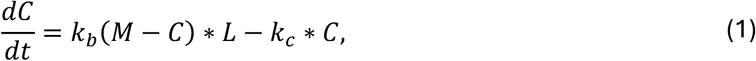

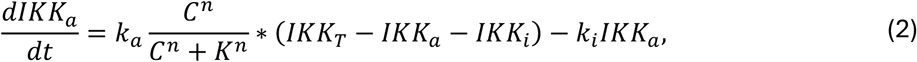

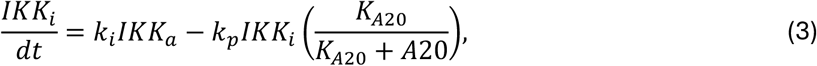

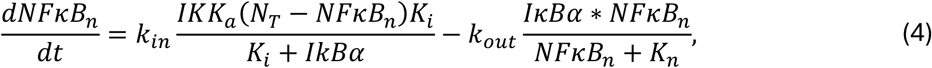

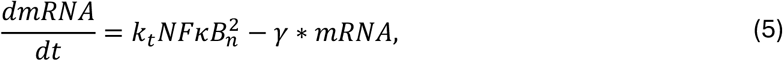

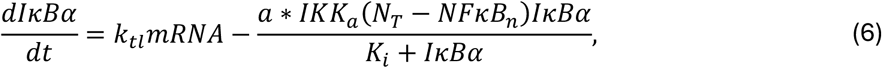

The activation function in our model depends on the total number of ligand-receptor complexes after stimulation. However, one can simplify this by normalizing the number of receptor-ligand complexes to the total number of receptors, which we assume is constant for each cell along the course of the experiment. This is, in Eq. (1) we can write *C* ≡ *C*/*M*, with the number of total receptors being absorbed by the constant *K* ≡ *K*/*M* in the activation function. In this form, the Hill function will only depend on the fraction of activated receptors. On the other hand, the units of the ligand dose can be absorbed into the receptor activation rate *k*_*b*_ ≡ *k*_*b*_*L*, setting the maximum ligand dose to one. Together, these modifications reduce the parameter space in our model to only 4 parameters. For completeness, below we rewrite the first equation with the aforementioned changes:

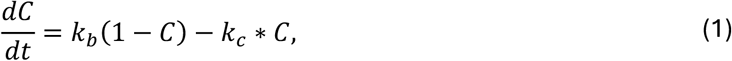

#### Model fitting and parameters

Our model uses 20 parameters. Table 1: 15 parameters were taken from the literature, one was measured (receptor expression) and the remaining four were fit. Table 2: using global optimization we fitted our model to experimental data to calculate the remaining 4 parameters.

**Table 1:** Default parameters for the minimal model in NF-κB. The number of receptors (M) for each pathway were measured and fitted to a Gamma distribution.

| # | Variable | Value | Unit | Description |
| --- | --- | --- | --- | --- |
| 1 | $k_{in}$ | .01 | 1/s | Tay et al. (28) |
| 2 | $k_{out}$ | .05 | 1/s | Tay et al. (28) |
| 3 | $k_a$ | $4 \cdot 10^{-3}$ | 1/s | Jensen et al. (42) |
| 4 | $k_i$ | $3 \cdot 10^{-3}$ | 1/s | Jensen et al. (42) |
| 5 | $k_p$ | $6 \cdot 10^{-4}$ | 1/s | Jensen et al. (42) |
| 6 | $k_t$ | $1.4 \cdot 10^{-7}$ | 1/(uM*s) | Ashall et al (20) |
| 7 | $k_{tl}$ | .5 | 1/s | Tay et al. (28) |
| 8 | $\gamma$ | $2.83 \cdot 10^{-4}$ | 1/s | Jensen et al. (42) |
| 9 | $a$ | 0.0175 | 1/(uM*s) | Jensen et al. (42) |
| 10 | $ka_{20}$ | 0.018 | uM | Jensen et al. (42) |
| 11 | $IKK_T$ | 2 | uM | Jensen et al. (42) |
| 12 | $K_n$ | .029 | uM | Jensen et al. (42) |
| 13 | $K_i$ | .035 | uM | Jensen et al. (42) |
| 14 | $N_T$ | 1 | uM | Jensen et al. (42) |
| 15 | $A_{20}$ | 0.0026 | uM | Jensen et al. (42) |
| 16 | $M_{TNFR}$ | $\Gamma(4.32, 1035)$ | | Measured |
| | $M_{TLR2}$ | $\Gamma(2.23, 4914)$ | | Measured |
| | $M_{TL27}$ | $\Gamma(4.80, 4404)$ | | Measured |

**Table 2:** Parameters fitted to experimental data for the three receptor systems TNFR, TLR2 and TLR7.

| # | Variable | Value |  |  | Unit |
| --- | --- | --- | --- | --- | --- |
|  |  | TNF | TLR2 | TLR7 |  |
| 1 | $k_b$ | 96 | .023 | .0120 | 1/(uM*s) |
| **2** | $k_c$ | .2 | .003 | .014 | 1/s |
| **3** | $n$ | 1.21 | 1.83 | 10 | |
| **4** | $K$ | 18.16 | 1.35 | 205.32 | |

Interestingly, the dissociation constant *k*_*d*_ = *k*_*c*_/ *k*_*b*_ exhibited by TNFR and TLR7 are close to those reported in the literature (*44*–*46*). For TLR2 we could not find a direct measurement reported yet.

