## supplementary information for "Accurate processing of ultra-short immune signals by single macrophages"

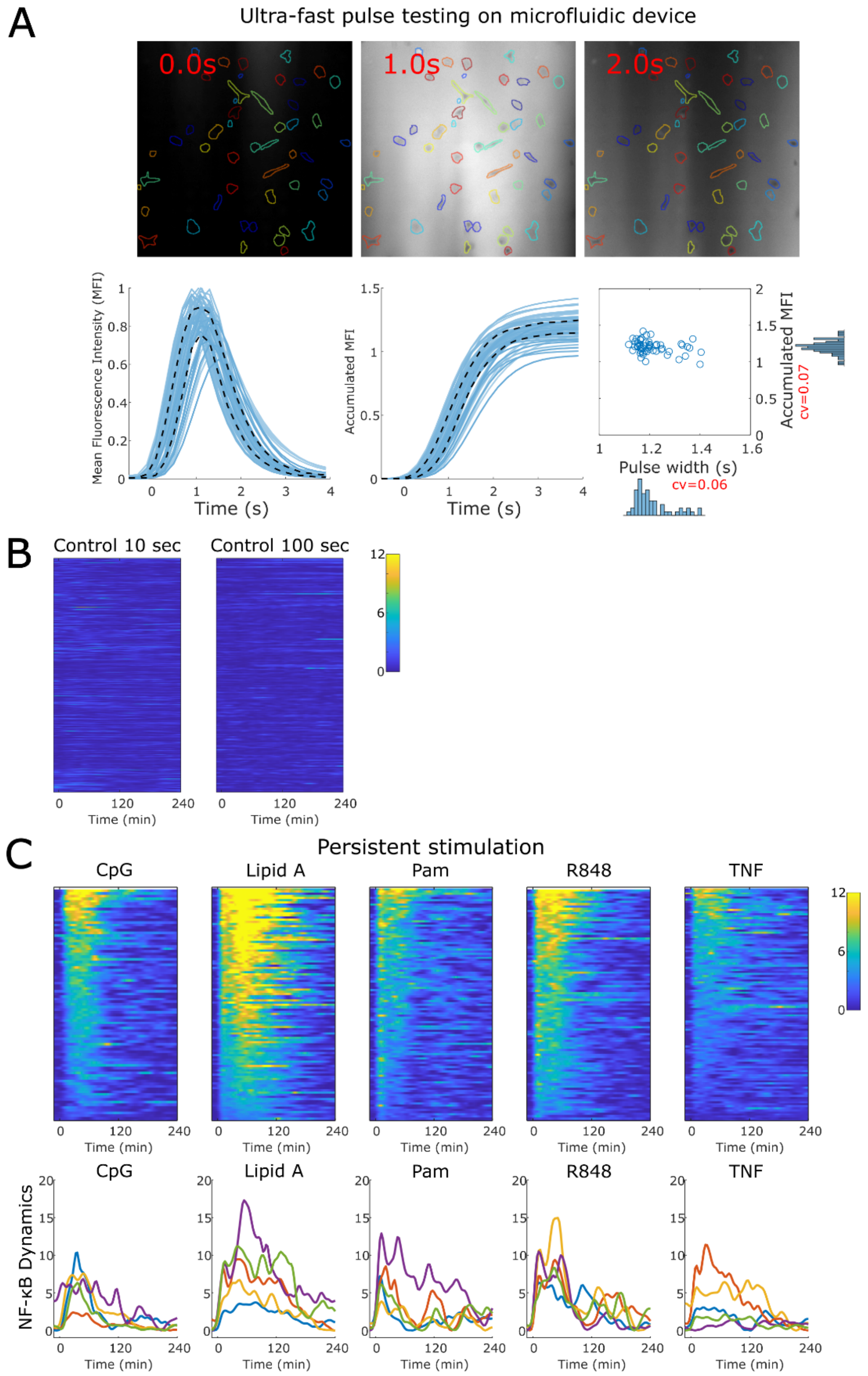


**Supplemental Figure 1:** A) Cy5 fluorescent signal for the shortest pulse within the microfluidic chamber. Fluorescence images were taken every 50 ms. We performed single-cell segmentation to measure the mean fluorescence intensity (MFI) on individual cells at each time point, as well as the accumulated MFI over 4 minutes, which reflects the total amount of effective medium that passes over a cell's area. Distribution of the pulse width as well as the total accumulated MFI on individual cells show very low variability, with a coefficient of variation around 0.06 and 0.07, respectively. From our measurements using flow cytometry, the receptor expression in macrophages varies by up to 0.5 around the mean. Since the variation in ligand concentration in our chambers is much lower, we consider receptor expression variability to be the main contributor to NF-κB heterogeneity, rather than ligand variation. B) Heatmap showing the responses of control cells exposed to 10-second and 100-second flows of fresh media in the microfluidic device, with 434 and 201 cells tracked, respectively. C) Heatmap and 5 random NF-kB traces for persistent stimulation.

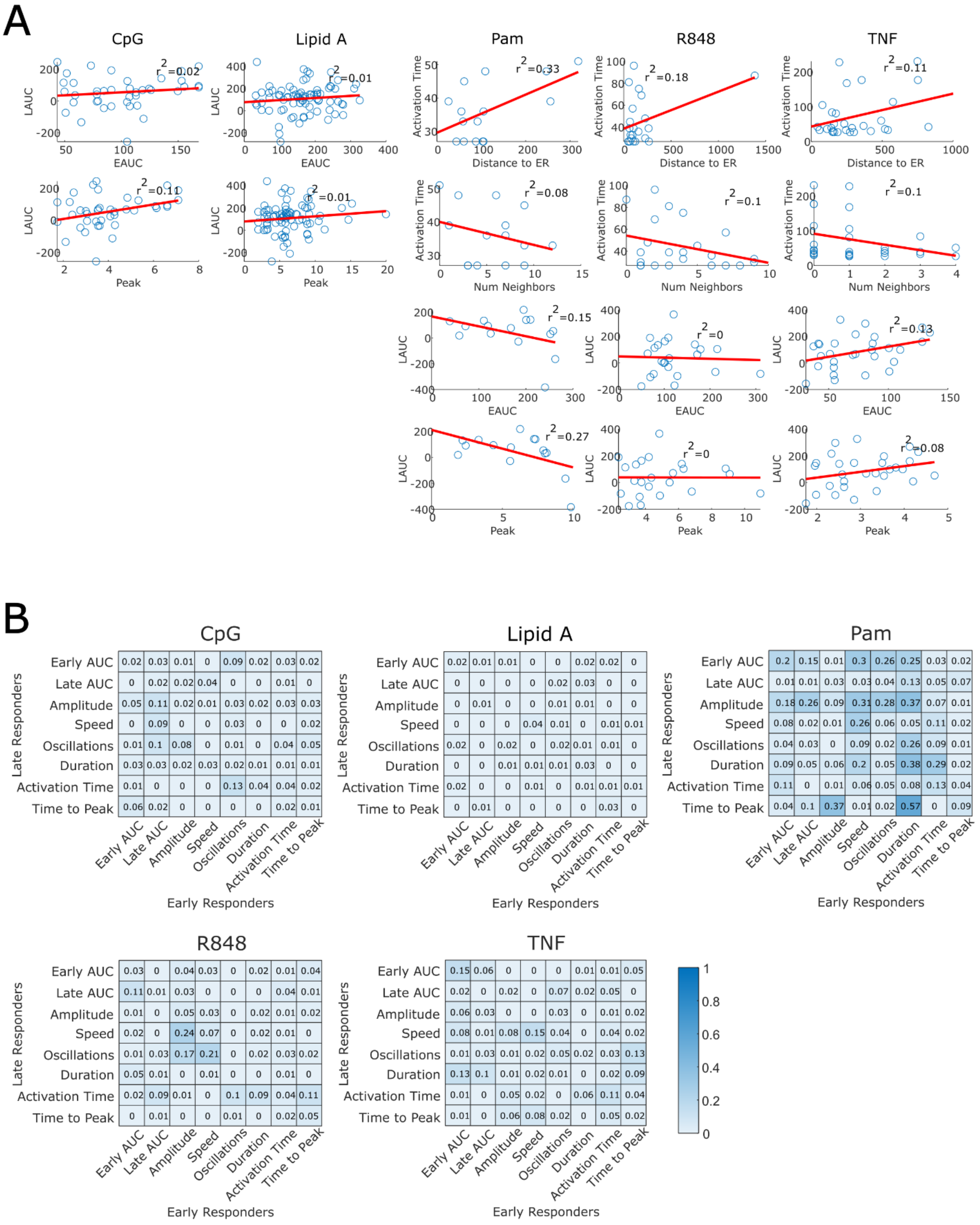


**Supplemental Figure 2:** A) Correlation between the late and early responders for individual cells and different metrics. B) Correlation matrices for the 8 NF-kB features between all late responders and their nearest early responder.

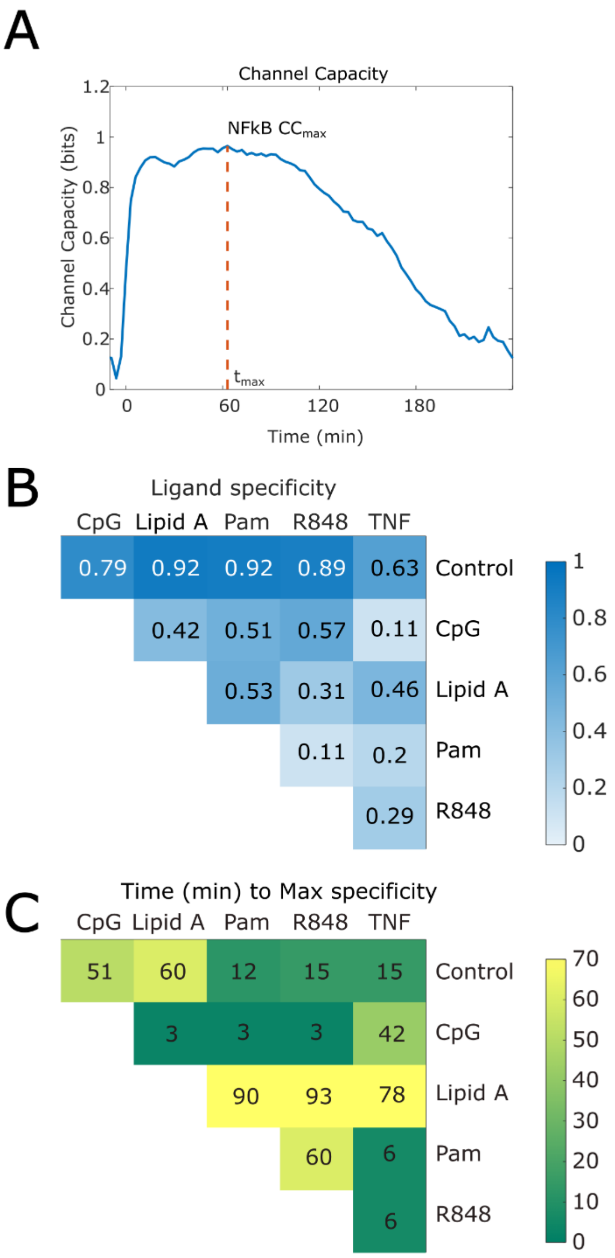


**Supplemental Figure 3:** A) Channel capacity, B) pair-wise mutual information and C) time to max specificity for macrophages under persistent stimulation.

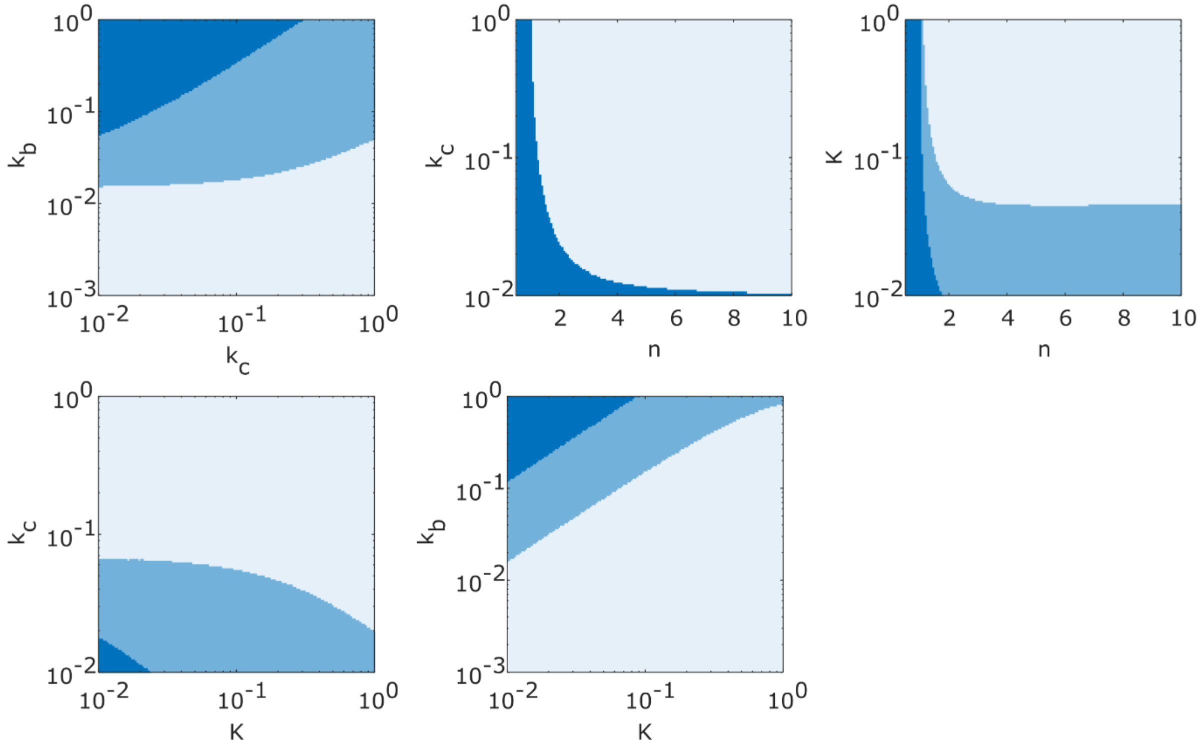


**Supplemental Figure 4:** Phase diagrams for any pair of parameters in our model. Blue colors represent the three different modes the NFkB strength can exhibit as a function of the pulse duration, according to the classification in main Figure 5C-D. Fix parameters are set to $k_b = .1\ 1/s, k_c = .03\ 1/s, K = .01, n = 2$.

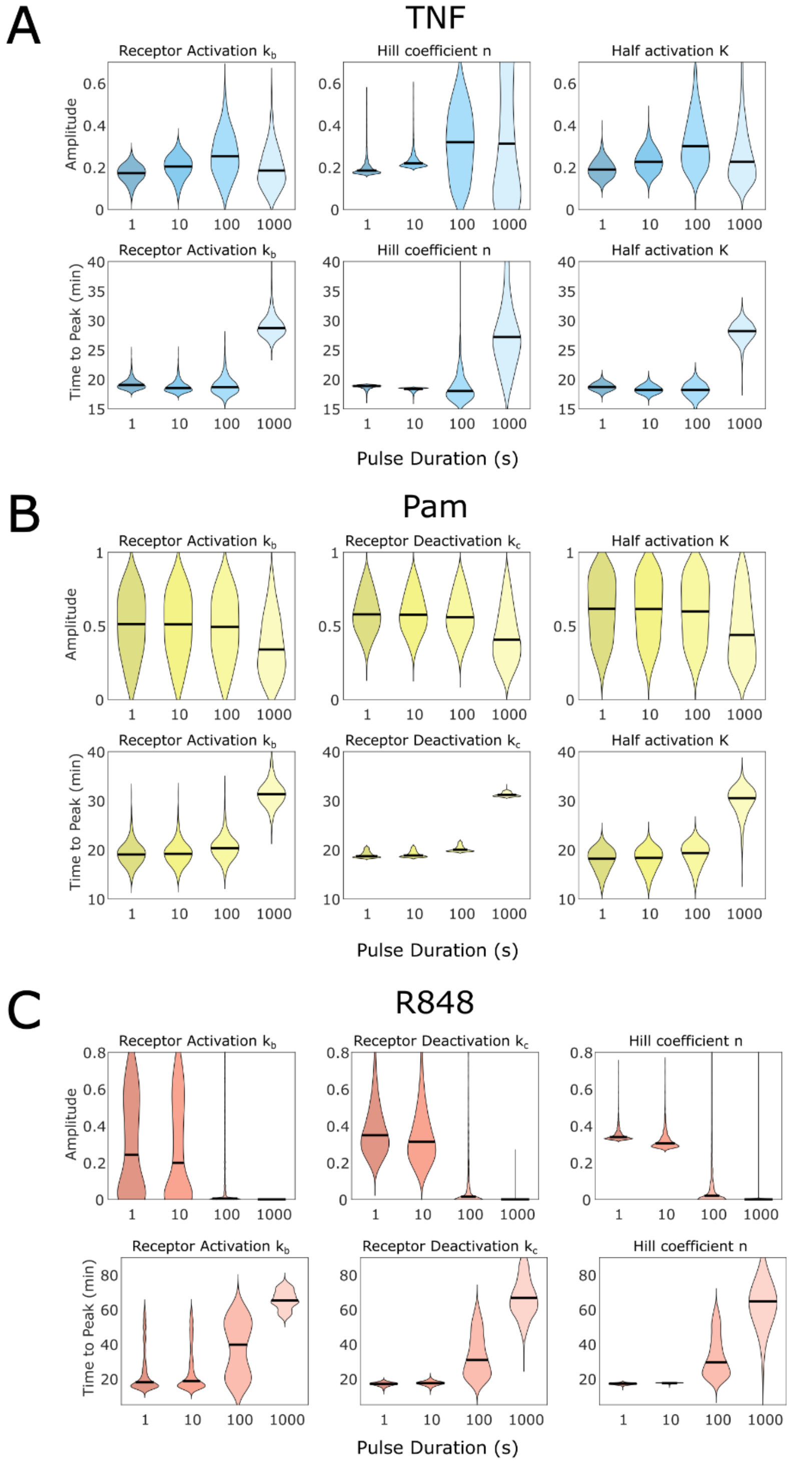


**Supplemental Figure 5:** Distribution of the amplitude and time to peak of NF-kB dynamics when individual parameter rates are varied for stimulation with A) TNF, B) PAM and C) R848. Each violin plot represents the distribution of 1000 cells. Black lines represent median values in the population.

**Supplemental Video 1:** Surface plot of the cy5 fluorescent signal for the shortest pulse within the microfluidic chamber, indicating signal dynamics within the microfluidic chamber. Cells within 75 micrometers from the side-walls of microfluidic chamber are excluded from the analysis to avoid signal duration variability at the edges of the chamber. As a result, the total signal pulse-width variability experienced by the analyzed cells is 0.07 (coefficient of variation). Also see Supplemental Figure 1.

**Supplemental Video 2:** Bone marrow derived primary macrophages in response to ultra-short stimulation with the 5 ligands we tested in this work.

**Supplemental Video 3:** Bone marrow derived primary macrophages in response to the cytokine TNF stimulation for four different pulse durations/amplitudes, keeping the area of stimulation constant.

**Supplemental Video 4:** Bone marrow derived primary macrophages in response to the pathogenic ligand Pam2CSK4 for four different pulse durations/amplitudes, keeping the area of stimulation constant.

**Supplemental Video 5:** Bone marrow derived primary macrophages in response to the small molecule R848 for four different pulse durations/amplitudes, keeping the area of stimulation constant.
